# Pleiotropic effects of *ebony* and *tan* on pigmentation and cuticular hydrocarbon composition in *Drosophila melanogaster*

**DOI:** 10.1101/538090

**Authors:** J. H. Massey, N. Akiyama, T. Bien, K. Dreisewerd, P. J. Wittkopp, J.Y. Yew, A. Takahashi

## Abstract

Pleiotropic genes are genes that affect more than one trait. For example, many genes required for pigmentation in the fruit fly *Drosophila melanogaster* also affect traits such as circadian rhythms, vision, and mating behavior. Here, we present evidence that two pigmentation genes, *ebony* and *tan*, which encode enzymes catalyzing reciprocal reactions in the melanin biosynthesis pathway, also affect cuticular hydrocarbon (CHC) composition in *D. melanogaster* females. More specifically, we report that *ebony* loss-of-function mutants have a CHC profile that is biased toward long (>25C) chain CHCs, whereas *tan* loss-of-function mutants have a CHC profile that is biased toward short (<25C) chain CHCs. Moreover, pharmacological inhibition of dopamine synthesis, a key step in the melanin synthesis pathway, reversed the changes in CHC composition seen in *ebony* mutants, making the CHC profiles similar to those seen in *tan* mutants. These observations suggest that genetic variation affecting *ebony* and/or *tan* activity might cause correlated changes in pigmentation and CHC composition in natural populations. We tested this possibility using the *Drosophila* Genetic Reference Panel (DGRP) and found that CHC composition covaried with pigmentation as well as levels of *ebony* and *tan* expression in newly eclosed adults in a manner consistent with the *ebony* and *tan* mutant phenotypes. These data suggest that the pleiotropic effects of *ebony* and *tan* might contribute to covariation of pigmentation and CHC profiles in *Drosophila*.

## 1 Introduction

When organisms adapt to novel environments, genetic changes often cause multiple traits to evolve. In some cases, organisms invading similar environments undergo similar shifts for suites of traits. In the threespine stickleback, for example, marine populations independently invading freshwater lake habitats have repeatedly evolved similar changes in defensive armor, behavior, and body shape (Walker and Bell, 2000; Schluter *et al.*, 2004; Wark *et al.* 2011). Such correlated evolution might result from (i) selection favoring a particular suite of traits (i.e. selection targeting multiple unlinked loci), (ii) selection favoring a trait that is genetically linked to genes affecting other traits, or (iii) selection favoring a trait that varies due to genetic variation at a pleiotropic gene affecting multiple traits. In the case of the threespine stickleback, genetic variation linked to a single major gene, Eda, has been found to explain correlated differences in these traits among populations (Albert *et al.*, 2008; Greenwood *et al.*, 2016), suggesting that pleiotropy has played a role. Studies in various other plant and animal species also support the hypothesis that pleiotropy contributes to the coevolution of correlated traits (e.g., McKay *et al.*, 2003; McLean *et al.*, 2011; Duveau and Felix 2012; Nagy *et al.* 2018).

In insects, genes determining body color are often pleiotropic. For example, in *Drosophila*, the *yellow* gene is required for the synthesis of black melanin and also affects mating behavior (Bastock, 1956; Drapeau *et al.*, 2003; Drapeau *et al.*, 2006). The genes *pale* and *Dopa-decarboxylase*, which encode enzymes that synthesize tyrosine-derived precursors for pigmentation, are also pleiotropic, affecting both body color and immunity (reviewed in Wittkopp and Beldade, 2009; Takahashi, 2013). In addition, prior work suggests that pigmentation genes might also affect cuticular hydrocarbon (CHC) profiles, which can affect desiccation (Gibbs, 1997; Gibbs, 1998; Foley and Telonis-Scott, 2011) and mate choice (reviewed in Yew and Chung, 2015). Specifically, a receptor for the tanning hormone *bursicon* and levels of the biogenic amine dopamine, which both affect cuticle pigmentation in *Drosophila melanogaster*, have been shown to influence CHC composition (Marican *et al.*, 2004;Wicker-Thomas and Hamann, 2008; Flaven-Pouchon *et al.* 2016).

Here, we test whether the *ebony* and *tan* genes of *D. melanogaster*, which are required for the synthesis of dark melanins and yellow sclerotins from dopamine, respectively, also affect CHC composition. The *ebony* gene encodes a protein that converts dopamine into N-β-alanyl dopamine (NBAD), and the *tan* gene encodes a protein that catalyzes the reverse reaction, converting NBAD back into dopamine (Figure 1A). We report that loss-of-function mutations in both *ebony* and *tan* altered CHC length composition relative to wild-type flies in opposing directions. These opposing effects on CHC length composition are consistent with *ebony* and *tan’s* opposing biochemical functions in dopamine metabolism (Figure 1A). Indeed, pharmacological inhibition of dopamine synthesis in *ebony* mutants caused a *tan*-like CHC length profile. To examine the possibility that variation in *ebony* and/or *tan* activity might cause correlated changes in pigmentation and CHC composition in a natural population, we used lines from the *Drosophila* Genetic Reference Panel (DGRP) to test for covariation between pigmentation and CHC composition. We found that CHC length composition covaried not only with pigmentation but also with levels of *ebony* and *tan* expression in a manner consistent with the mutant analyses. In the discussion, we compare our data to studies of clinal variation in CHC composition and pigmentation to determine whether the pleiotropic effects we see might have contributed to correlated evolution of these traits.

**Figure 1.**
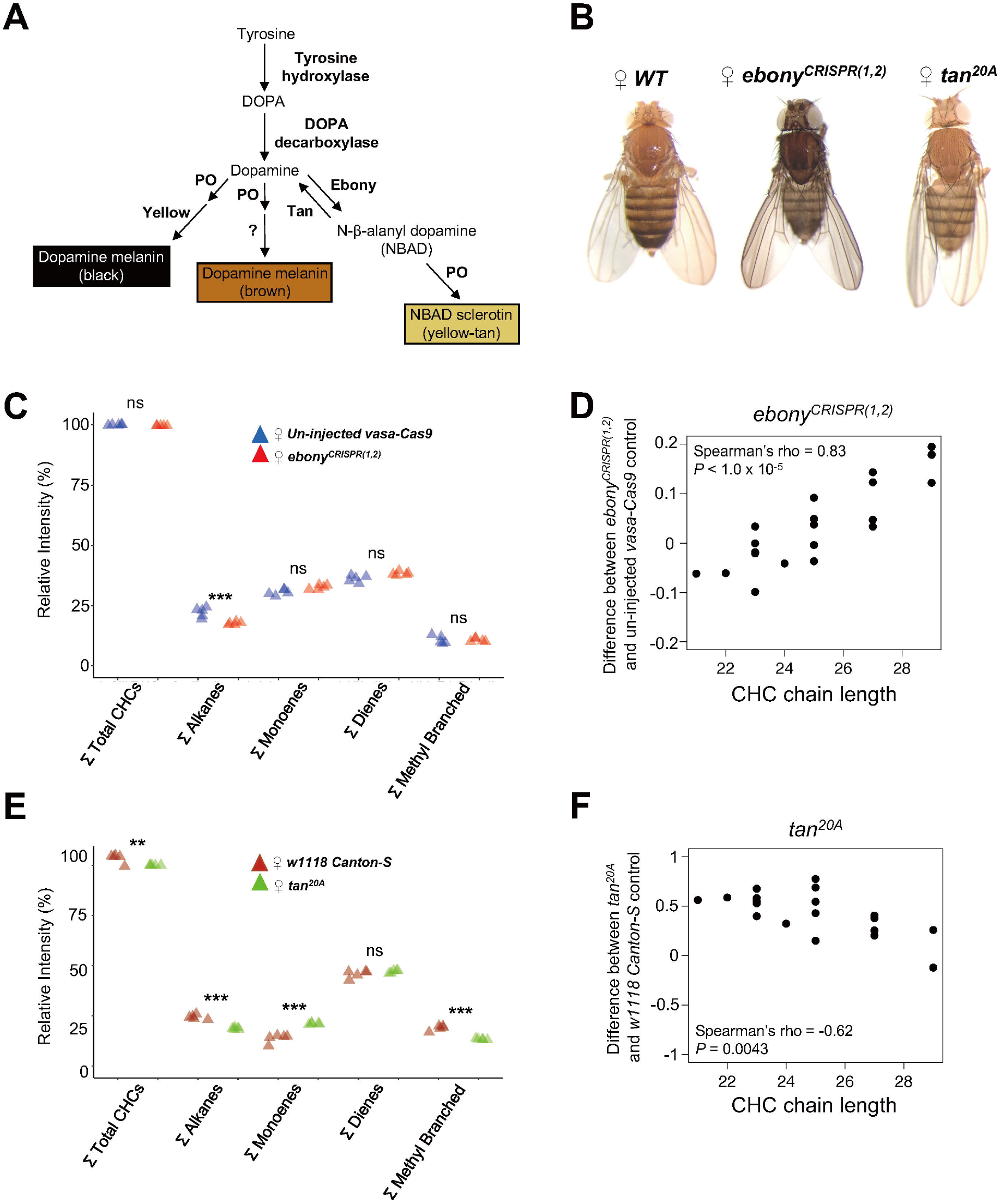
*ebony* and *tan* affect pigmentation and CHC composition in female *Drosophila* melanogaster. (**A**) Insect sclerotization and pigmentation synthesis pathway. Ebony converts dopamine into N-β-alanyl dopamine (NBAD) which is oxidized into yellow-colored NBAD sclerotin. Tan catalyzes the reverse reaction, converting NBAD back into dopamine that can be oxidized into black and brown melanins. (**B**) Photographs highlighting the effects of *ebony^CRISPR(1,2)^* (darker) and *tan^20A^* (lighter) on body pigmentation compared to the un-injected *vasa-Cas9* control line (*WT*). (**C**) Summary of *ebony^CRISPR(1,2)^* effects on total summed CHC classes relative to *un-injected vasa-Cas9* control females. (**D**) Difference in log-contrast of relative CHC intensity between *ebony^CRISPR(1,2)^* and *uninjected vasa-Cas9* control flies. (**E**) Summary of *tan^20A^* effects on total summed CHC classes relative to *w^1118^Canton-S* control females. For (D) and (E), each triangle represents a single replicate of CHCs extracted from five pooled individuals (N = 5 replicates per genotype). (**F**) Difference in log-contrast of relative CHC intensity between *tan^20A^* and *w^1118^ Canton-S* control flies. Results of Tukey HSD post-hoc tests following One-way ANOVA are shown: * P < 0.05, ** P < 0.01, *** P < 0.001.

## 2 Materials and Methods

### 2.1 Fly stocks and maintenance

The following lines were used: P excision line *tan^20A^* (True *et al.*, 2005) (courtesy of John True, Stony Brook University); the *UAS-ebony-RNAi* effector line was obtained from the Vienna Drosophila Resource Centre (Dietzl *et al.*, 2007, KK106278); *dsx^GAL4^* (Rideout *et al.*, 2010) (courtesy of Stephen Goodwin, Oxford University); *OK72-GAL4* (Ferveur *et al.*, 1997) (courtesy of Scott Pletcher, University of Michigan); *pannier-GAL4* (Calleja *et al.* 2000) was obtained from the Bloomington Drosophila Stock Center (BDSC 3039); *vasa-Cas9* (Gratz *et al.*, 2014, BDSC 51324) (courtesy of Rainbow Transgenics Inc.). All flies were grown at 23°C with a 12 h light-dark cycle on standard corn-meal fly medium.

#### 2.1.1 DGRP stocks

The following inbred *D. melanogaster* lines from the DGRP (Ayroles *et al.* 2009; Mackay *et al.* 2012; Huang *et al.* 2014) were used in this study: RAL-208, RAL-303, RAL-324, RAL-335, RAL-357, RAL-358, RAL-360, RAL-365, RAL-380, RAL-399, RAL-517, RAL-555, RAL-705, RAL-707, RAL-732, RAL-774, RAL-786, RAL-799, RAL-820, RAL-852, RAL-714, RAL-437, RAL-861 and RAL-892. These lines consist of the set of 20 lines used in Miyagi *et al.* (2015) and additional 3 dark lines (RAL-714, RAL-437, and RAL-861), which were added to avoid line specific effects from a limited number of dark lines. All flies were grown at 25°C with a 12 h light-dark cycle on standard corn-meal fly medium.

### 2.2 Generation of *ebony* CRISPR lines

New loss-of-function *ebony* mutants were constructed by synthesizing two single guide RNAs (gRNA), using a MEGAscript T7 Transcription Kit (Invitrogen), following the PCR-based protocol from Bassett *et al.* (2014), that target *ebony*’s first coding exon and co-injecting these at a total concentration of 100 ng/μL into embryos of a *D. melanogaster vasa-Cas9* line (Gratz *et al.*, 2014; BDSC 51324) (Supplementary Figure S1). These gRNAs were previously found to generate a high level of heritable germline transformants (Ren *et al.*, 2014; Supplementary Figure 1). We screened for germline transformants based on body pigmentation and confirmed via Sanger sequencing three unique *ebony* loss-of-function alleles, *ebony^CRISPR(1,2)^* containing a 55 bp deletion, and *ebony^CRISPR(3)^* and *ebony^CRISPR(4)^* each containing an in-frame 3 bp deletion (Supplementary Figure S1). Each deletion caused flies to develop dark body pigmentation, indicating loss of Ebony activity (Figure 1B, Supplementary Figure S2A).

### 2.3 CHC extraction and measurements

For Figures 1 and 2 and Supplementary Figures S2–5, CHCs were extracted and analyzed as described below (CHC names and formulas are summarized in Supplementary Table S1). For the analyses using the DGRP (Figures 3 and 4, Supplementary Figure S6), all CHC data for females were obtained from Dembeck *et al.* (2015b); however, in the case of GC/MS peaks composed of more than two combined CHC components that differed in CHC chain length, the non-branched CHC chain length was used. Also, CHCs that were not detected in all strains were removed from the analyses.

**Figure 2.**
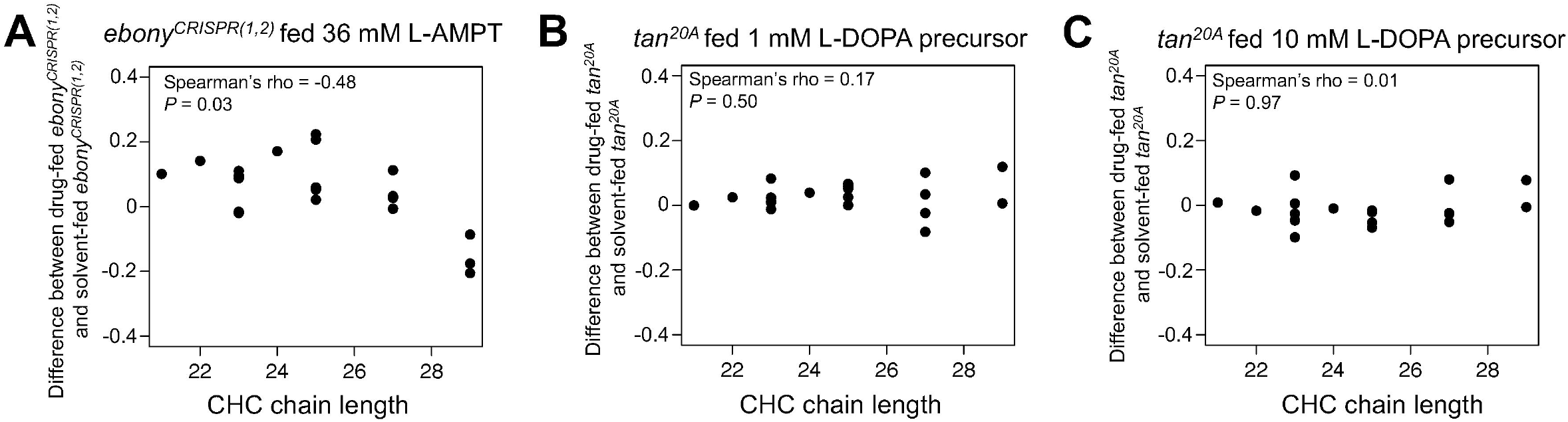
Effects of pharmacological treatments on CHC lengthening in *ebony^CRISPR(1,2)^* and *tan^20A^* mutants. **(A)** Difference in log-contrast of relative CHC intensity between *ebony^CRISPR(1,2)^* females fed 36mM alpha methyl tyrosine (L-AMPT) and *ebony^CRISPR(1,2^* females fed a solvent control. **(B)** Difference in log-contrast relative of CHC intensity between *tan^20A^* females fed 1 mM methyl L-DOPA hydrochloride (L-DOPA precursor) and *tan^20A^* females fed a solvent control. **(C)** Difference in log-contrast of relative CHC intensity between *tan^20A^* females fed 10 mM L-DOPA precursor and *tan^20A^* females fed a solvent control.

**Figure 3.**
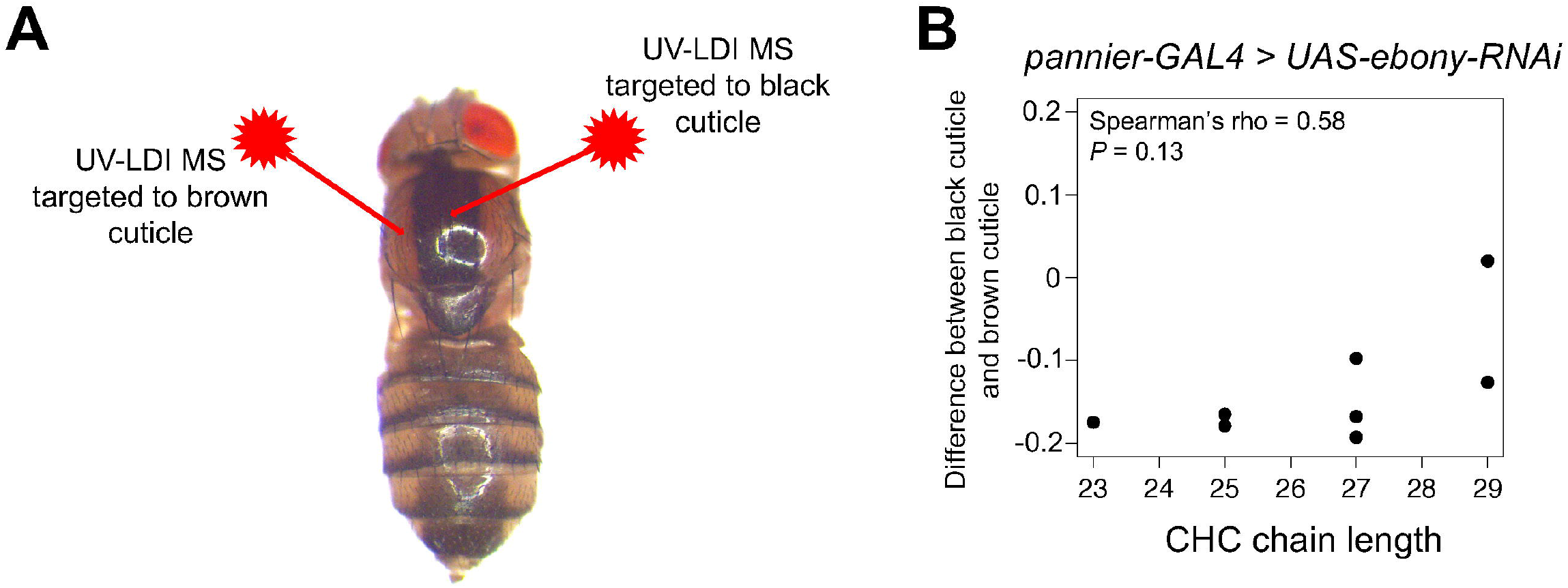
UV laser desorption/ionization mass spectrometry (UV-LDI MS) did not detect differences in short versus long CHCs between lightly and darkly pigmented cuticle. Female *pannier-GAL4* flies were crossed to *UAS-ebony-RNAi* males to generate flies with a dark, heavily melanized stripe down the dorsal midline. **(A)** The UV-LDI MS lasers were targeted to light brown or dark black cuticle within the same fly (N = 3 biological replicates). **(B)** Difference in relative CHC intensity between black and brown cuticle.

**Figure 4.**
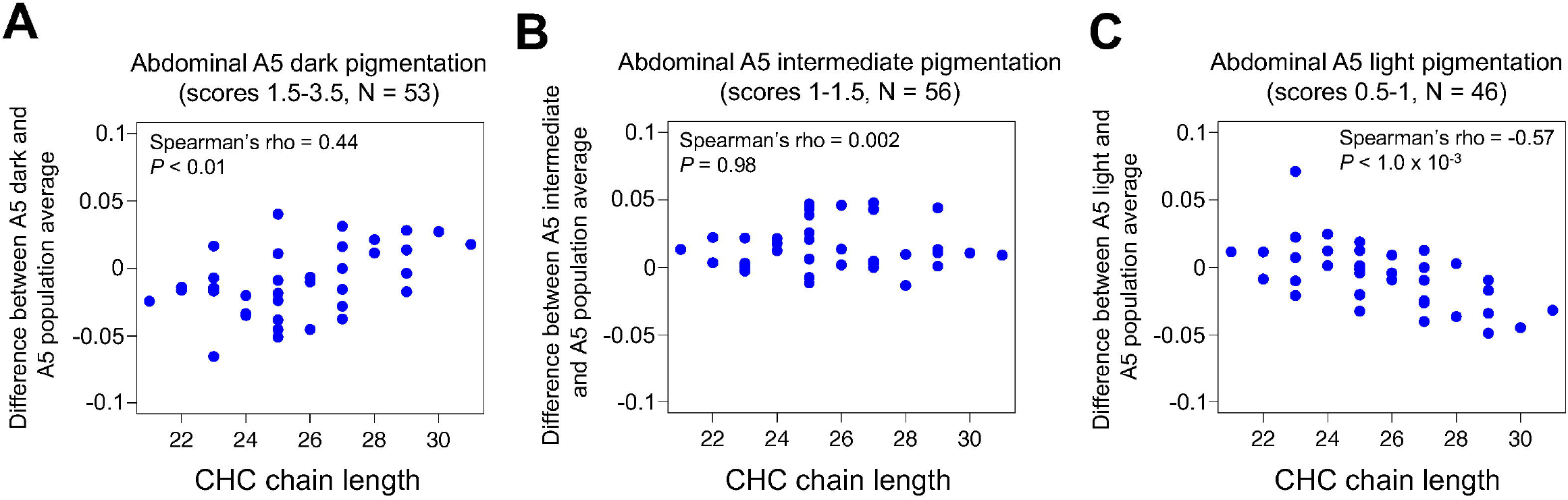
Abdominal pigmentation co-varies with CHC length profiles in the *Drosophila* Genetic Reference Panel (DGRP) Pigmentation scores and CHC data were obtained from Dembeck *et al.* (2015a,b). **(A)** Difference in log-contrast of relative CHC intensity between DGRP females with darkly-pigmented 5^th^ abdominal tergites (A5) (1.5 < score, N = 53) and the 155 line average. **(B)** Difference in log-contrast of relative CHC intensity between DGRP females with intermediately-pigmented A5 (1 < score ≤ 1.5, N = 56) and the 155 line average. **(C)** Difference in log-contrast of relative CHC intensity between DGRP females with lightly-pigmented A5 (score ≤ 1, N = 49) and the 155 line average.

#### 2.3.1 Extraction

For each experiment, five replicate CHC samples of virgin female flies were prepared for each genotype or pharmacological treatment group. All *ebony* and *tan* mutant CHC extractions were performed on 3–4 d old virgin females. For pharmacological experiments, 1–2 d old virgin females were treated for 4 d prior to CHC extraction. For GAL4/UAS experiments, virgin females were tested at 10–12 d. For each sample, 5 flies were placed in a single glass vial (Wheaton 224740 E−C Clear Glass Sample Vials) on ice. 120 μL of hexane (Sigma Aldrich, St Louis, MO, USA) spiked with 10 μg/mL of hexacosane (Sigma Aldrich) was added to each vial and sealed with a cap. Vials were incubated at room temperature for 20 mins. 100 μL of the cuticular extract was removed, transferred into a clean vial (Wheaton 0.25 mL with low volume insert), and stored at −20°C.

#### 2.3.2 GC/MS analysis

Gas chromatography mass spectrometry (GC/MS) analysis was performed on a 7820A GC system equipped with a 5975 Mass Selective Detector (Agilent Technologies, Inc., Santa Clara, CA, USA) and a HP-5ms column ((5%-Phenyl)-methylpolysiloxane, 30 m length, 250 μm ID, 0.25 μm film thickness; Agilent Technologies, Inc.). Electron ionization (EI) energy was set at 70 eV. One microliter of the sample was injected in splitless mode and analyzed with helium flow at 1 mL/ min. The following parameters were used: column was set at 40°C for 3 min, increased to 200°C at a rate of 35°C/min, then increased to 280°C at a rate of 20°C/min for 15 min. The MS was set to detect from *m/z* 33 to 500. Chromatograms and spectra were analyzed using MSD ChemStation (Agilent Technologies, Inc.). CHCs were identified on the basis of retention time and EI fragmentation pattern. The relative abundance for each CHC signal was calculated by normalizing the area under each CHC peak to the area of the hexacosane signal. To eliminate multicollinearity among sample peak amounts, a log-contrast transformation was applied to the resulting proportional values, using nC27 as the denominator (Yew *et al.*, 2011; Blow and Allen, 1998):

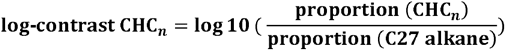

To determine the relative change in CHC length between two genotypes, experimental groups, or groups of DGRP strains, the difference in relative intensity of individual CHC intensities of each group was calculated:

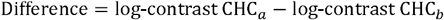

These values were then plotted against CHC chain length.

### 2.4 Ultraviolet laser desorption ionization mass spectrometry (UV-LDI MS)

For intact fly analysis, individual animals were attached to a glass cover slip using adhesive pads (G304, Plano, Wetzlar, Germany). The cover slips were mounted on a custom-milled sample holder containing a rectangular, 1.8 mm deep well. Sample height was adjusted by choosing a stack of 0.2 mm-thick adhesive pads (G3347, Plano). Mass spectra were generated using a prototype orthogonal-extracting mass spectrometer (oTOF-MS) as described previously (Yew *et al.* 2011). The oTOF-MS was equipped with a modified oMALDI2 ion source (AB Sciex, Concord, Canada) and an N_2_ laser (λ = 337 nm) operated at a pulse repetition rate of 30 Hz. N_2_ was used as buffer gas at p = 2 mbar. This elevated pressure is critical to achieve an efficient collisional cooling environment for generation of weakly-bound [M + K]+ ions that constituted the major molecular ion species. Before starting the actual measurements, external mass calibration was achieved with red phosphorus, resulting in a mass accuracy of approximately 25 ppm. Approximately 900 laser shots were placed at one position to achieve a mass spectrum (30 s @30 Hz). All spectra were acquired in positive ion mode and processed using MS Analyst software (Analyst QS 2.0, AB Sciex, Concord, Canada).

### 2.5 Pharmacology Experiments

For pharmacological treatments, standard corn-meal fly medium was liquefied and cooled to ca. 60°C before the addition of each respective drug or solvent control. Ten 1–2 d old virgin females were placed in the vials for 4 d. To inhibit tyrosine hydroxylase activity, we prepared a 36 mM alpha methyl tyrosine (L-AMPT) (Sigma Aldrich) diet. The pH of the solution was adjusted with concentrated HCl until the drug dissolved. A solvent control diet solution was prepared using identical procedures. For the dopamine treatments, 1 mM and 10 mM L-dopa precursor (Methyl L-DOPA hydrochloride) (Sigma Aldrich) were dissolved in water before adding to liquefied fly media.

### 2.6 RNA extraction

Female virgin flies were collected within 1 h of eclosion, and the heads were removed in RNAlater (Ambion) to separate the effect from transcripts in non-epidermal head tissues. The remaining headless body samples were stored in RNAlater at −80°C until use. Three body samples from each line were placed in a 2 mL microtube with 400 μL TRIzol Reagent (Thermo Fisher Scientific, Tokyo, Japan) and an equivalent volume of 1.2 mm zirconia silica beads (Bio Medical Science). After shaking the tube at 3,200 rpm for 2 min using a Beads Crusher μT-12 (TAITEC, Koshigaya, Japan), 160 μl chloroform was added and mixed thoroughly. Total RNA in the aqueous phase was subsequently purified using silica-gel (Wakocil 5SIL, Wako, Osaka, Japan) based on the method of Boom *et al.* (1990) and was quantified using a Nanodrop 2000c spectrophotometer (Thermo Fisher Scientific).

### 2.7 Quantitative real-time PCR (qRT-PCR)

First strand cDNA was synthesized from 1 μg total RNA by using a PrimeScript RT Reagent Kit with gDNA Eraser (Takara Bio, Kusatsu, Japan). qRT-PCR was performed in a 25 μl reaction volume with SYBR Premix Ex Taq II Tli RNaseH Plus (Takara Bio) on a Thermal Cycler Dice TP800 (Takara Bio). Primer pairs used for RT-qPCR were *ebony*. 5’–CTTAGTGTGAAACGGCCACAG–3’ and 5’-GCAGCGAACCCATCTTGAA-3’; *tan*: 5’-GTTGAGGGGCTTCGATAAGA-3’ and 5’–GTCCTCCGGAAAGATCCTG–3’; *Act57B*: 5’–CGTGTCATCCTTGGTTCGAGA–3’ and 5’–ACCGCGAGCGATTAACAAGTG–3’; *Rp49*: 5’–TCGGATCGATATGCTAAGCTG–3’ and 5’–TCGATCCGTAACCGATGTTG–3’. *Act57B* and *Rp49* were used as internal control. Two replicate PCR reactions were performed for each cDNA sample and three biological replicates were obtained for each line.

### 2.8 Grouping DGRP lines based on pigmentation scores and *ebony/tan* expression levels

The DGRP lines (N = 155) with both pigmentation scores in Dembeck *et al.* (2015a) and CHC profiles in Dembeck *et al.* (2015b) were grouped into dark, intermediate, and light pigmentation lines using the pigmentation scores of the abdominal tergites from Dembeck *et al.* (2015a). The scores ranged from 0 for no dark pigmentation to 4 for 100% dark pigmentation in increments of 0.5, and were averaged across 10 individuals per line. Pigmentation grouping was done based on the score delimitations that split the lines most evenly into three groups. For the 5^th^ tergite (A5), lines were categorized into following groups: dark (1.5 < score, N = 53), intermediate (1 < score ≤ 1.5, N = 56), and light (score ≤ 1, N = 49). For the 6^th^ tergite (A6), lines were categorized into following groups: dark (3 < score, N = 51), intermediate (2 < score ≤ 3, N = 55), light (score ≤ 2, N = 49).

The 23 DGRP lines with varying *ebony* and *tan* expression levels were grouped into low, intermediate, and high expression lines using the qRT-PCR data. Since the normalized quantities are continuous values, grouping was done based on standard deviations (SD). For the *ebony* expression, lines were categorized into following groups: low (expression < mean - 0.5SD, N = 6), intermediate (mean - 0.5SD ≤ expression ≤ mean + 0.5SD, N = 9), and high (mean + 0.5SD < expression, N = 8). For the *tan* expression, lines were categorized into following groups: low (expression < mean - 0.5SD, N = 10), intermediate (mean - 0.5SD ≤ expression ≤ mean + 0.5SD, N = 7), and high (mean + 0.5SD < expression, N ≤ 6).

### 2.9 Statistics

All statistical tests were performed in R for Mac version 3.3.3 (R Core Team 2018) using one-way ANOVAs to test for statistically significant effects between more than two groups and post-hoc Tukey HSD tests for multiple pairwise comparisons. We used Spearman’s rank correlation coefficient ρ to test for the significance of the association. All pairwise tests were two-tailed, and the level of significance was set as *α* = 0.05.

## 3 Results

### 3.1 Loss-of-function mutations in *ebony* and *tan* have reciprocal effects on CHC length profiles

To determine whether the *ebony* gene affects cuticular hydrocarbons (CHCs), we created three new *ebony*-mutant alleles via CRISPR/Cas9 gene editing. One allele, *ebony^CRISP(1,2)^* contained a 55 bp deletion that caused a frame-shift in *ebony*’s coding sequence (Supplementary Figure S1C). Flies homozygous for this *ebony^CRISPR(1,2^* allele showed dark body pigmentation similar to that described previously for loss-of-function *ebony* mutants (Bridges and Morgan, 1923) (Figure 1B). We measured CHC profiles in 3–4 d old *ebony^CRISPR(1,2)^* virgin females using gas chromatography (GC/MS) and found that *ebony^CRISPR(1,2)^* flies showed lower levels of total alkanes relative to 3–4 d old virgin females from the strain the guide RNAs were injected into (i.e., un-injected *vasa-Cas9*)(Figure 1C, One-way ANOVA: F_9,40_ = 4494, *P* < 2.0 x 10^-16^; post-hoc Tukey HSD was significant for alkanes: *P* < 1.0 x 10^-5^).

We then tested whether *ebony^CRISPR(1,2)^* females had different proportions of individual CHCs. We calculated the average difference in individual log-contrast transformed CHC relative intensities (see Materials and Methods) between *ebony^CRISPR(1,2)^* flies and un-injected *vasa-Cas9* control flies and plotted these values against CHC chain length (varying from 21 carbons (C) to 29C) (Figure 1D, Supplementary Table S1). We found that *ebony^CRISPR(1,2)^* flies tended to show lower levels of short chain CHCs (<25C) and higher levels of long chain CHCs (>25C), suggesting that disrupting the function of ebony causes a CHC lengthening effect (Figure 1D, Spearman’s ρ = 0.83, *P* < 1.0 x 10^-5^).

The two other *ebony* alleles generated using CRISPR/Cas9 gene editing (*ebony^CRISPR(3)^* and *ebony^CRISPR(4)^*) each had a single 3 bp in-frame deletion in the first coding exon (Supplementary Figure S1D,E), suggesting that they might have less severe effects on Ebony activity than the *ebony^CRISPR(1,2)^* allele containing a 55 bp deletion causing a frame-shift. Consistent with this prediction, these ebony mutants also showed darker body pigmentation than wild-type flies (Supplementary Figure S2A), but did not show any bias toward longer CHCs (Supplementary Figure S2B,C, *ebony^CRISPR(3)^*: Spearman’s ρ = 0.22, *P* = 0.34; *ebony^CRISPR(4)^*: Spearman’s ρ = 0.07, *P* = 0.78).

To better understand the effects of reduced *ebony* expression on CHCs, we knocked down *ebony* expression in specific cell types using ebony-RNAi (Dietzl *et al.*, 2007). First, we drove expression of ebony-RNAi with the *dsx^GAL4^* driver (Rideout *et al.*, 2010), which causes RNAi expression in the cuticle, fat body, CNS, and oenocytes among other tissues. We observed darker pigmentation in *dsx^GAL4^ > UAS-ebony-RNAi* flies than control flies (data not shown), suggesting that the ebony-RNAi effectively targeted and knocked down *ebony* expression. These *dsx^GAL4^ > UAS-ebony-RNAi* flies also showed a pattern of CHC lengthening similar to the *ebony^CRISPR(1,2^* mutants when compared to to *dsx^GAL4^* / + control flies but not when compared to *UAS-ebony-RNAi* / + control flies. This result might be due to leaky *UAS-ebony-RNAi* expression in the latter control flies that makes their profiles more similar to those of *dsx^GAL4^> UAS-ebony-RNAi* flies (Supplementary Figure S3A, B, relative to *dsx^GAL4^ / +* control: Spearman’s ρ = 0.58, *P* < 0.007; relative to *UAS-ebony-RNAi* /+ control: Spearman’s ρ = 0.19, *P* = 0.42).

We hypothesized that the effect on CHCs might be due to reducing *ebony* expression specifically in oenocytes because these cells synthesize many CHC precursor compounds (Wigglesworth, 1970). Therefore, we drove expression of ebony-RNAi using the *OK72-GAL4* driver that is also expressed in oenocytes (Ferveur *et al.*, 1997). These flies showed no significant difference in CHC length profiles (Supplementary Figure S3C, Spearman’s ρ = −0.01, *P* = 0.96), suggesting that *ebony* expression in non-oenocyte tissues expressing *doublesex* affects the overall length proportion of CHCs.

Next, we asked whether loss-of-function mutations in the *tan* gene also affect CHC composition. Specifically, we examined CHC composition in 3–4 d old virgin females carrying a *tan^20A^* null allele, which contains an imprecise P-element excision that results in a 953 bp deletion that includes the presumptive promoter region (True *et al.*, 2005). Because *tan* encodes a protein that catalyzes the reverse of the reaction catalyzed by Ebony (Figure 1A), we predicted that *tan* mutants might show the opposite effects on CHC composition. Similar to the *ebony^CRISPR(1,2)^* mutants, *tan^20A^* females showed differences in the overall abundance of alkanes, but also total CHCs, monoenes, and methyl branched CHCs (Figure 1E, One-way ANOVA: F_9,40_ = 3586, *P* < 2.0 x 10^-16^; post-hoc Tukey HSD was significant for total summed CHCs: P < 0.01, total summed alkanes: *P* < 0.001, total summed monoenes: P < 0.001, and total summed methyl branched: *P* < 0.001). More importantly, *tan^20A^* females tended to show higher levels of short chain CHCs relative to long chain CHCs when compared to *w^1118^ Canton-S (CS)* control flies, as predicted (Figure 1F, Spearman’s ρ = −0.62, *P* = 0.0043). Together, these results suggest that *ebony* and *tan* have reciprocal effects on both pigmentation synthesis (reviewed in True, 2003 and True, 2005) and CHC length profiles. We note that this conclusion contradicts Wicker-Thomas and Hamann (2008)’s report that CHC profiles were similar in *ebony* or *tan* loss-of-function mutants and wild-type flies; however, the *ebony* and *tan* alleles used in this prior work might not have been nulls.

### 3.2 Pharmacological inhibition of tyrosine hydroxylase activity reverses the CHC lengthening effect in *ebony^CRISPR(1,2)^* flies

We hypothesized that *ebony* and *tan* might have reciprocal effects on CHC length profiles because of their effects on dopamine metabolism. For example, because *ebony* encodes a protein that converts dopamine into NBAD (Figure 1A), we hypothesized that loss-of-function *ebony* mutants might accumulate dopamine (as reported in Hodgetts and Konopka, 1973) and that this dopamine might be shunted into other pathways, possibly affecting CHC lengthening. To explore this hypothesis, we fed 1-2 d old adult female *ebony^CRISPR(1,2)^* flies a tyrosine hydroxylase inhibitor, alpha methyl tyrosine (L-AMPT), for four days to determine whether inhibiting dopamine synthesis would reverse the CHC lengthening pattern we observed in *ebony^CRISPR(1,2)^* flies. Relative to *ebony^CRISPR(1,2)^* solvent-fed control flies, *ebony^CRISPR(1,2)^* flies fed 36 mM L-AMPT did indeed reverse the CHC lengthening pattern we observed in *ebony^CRISPR(1,2)^* flies, resulting in a shortening of CHCs similar to that observed in *tan^20A^* flies (Figure 2A, Spearman’s ρ = −0.48, *P* = 0.03). Feeding 1–2 d old adult flies L-AMPT did not, however, affect body pigmentation (data not shown), consistent with body pigmentation being determined prior to and soon after eclosion (Hovemann *et al.*, 1998). We also fed *ebony^CRISPR(4)^* flies a 36 mM dose of L-AMPT to see if we could induce CHC shortening in an *ebony* mutant with unchanged CHC length composition. Similar to *ebony^CRISPR(1,2)^* fed flies, we detected a significant negative correlation when comparing *ebony^CRISPR(4)^* fed flies to an *ebony^CRISPR(4)^* solvent-fed control (Supplementary Figure S4, Spearman’s ρ = −0.57, *P* = 0.009).

We next hypothesized that *tan^20A^* flies might have lower levels of circulating dopamine, because *tan* encodes a protein that converts NBAD back into dopamine (Figure 1A). To determine whether elevating dopamine levels in *tan* mutants would affect CHCs, we fed *tan^20A^* females a dopamine precursor, methyl L-DOPA hydrochloride (L-DOPA precursor), to see if elevating dopamine levels could reverse the CHC shortening pattern we observed in *tan^20A^* flies; however, neither the 1 mM nor 10 mM L-DOPA precursor treatments seemed to affect CHC length profiles when compared to *tan^20A^* solvent-fed control flies (Figure 2B, C, Spearman’s ρ = 0.17, *P*= 0.50; Spearman’s ρ = 0.01, *P* = 0.97, respectively). We also fed *tan^20A^* flies a higher 100 mM dose of the L-DOPA precursor, but all of these flies died before CHC extraction; these flies also showed darker cuticle pigmentation consistent with elevated dopamine. Finally, we fed 1 mM and 10 mM doses of L-DOPA precursor to wild-type (*w^1118^ CS*) females to see if we could induce CHC lengthening in a wild-type genetic background; instead, we observed a slight CHC shortening effect for the 1 mM dose and no effect for the 10 mM dose (Supplementary Figure S5, Spearman’s ρ = −0.52, *P* = 0.02; Spearman’s ρ = −0.36, *P* = 0.12, respectively). Together, these results indicate that inhibiting tyrosine hydroxylase activity in *ebony* mutants causes a CHC shortening effect like that observed in *tan^20A^* flies; however, increasing dopamine levels through feeding does not cause a CHC lengthening effect.

### 3.3 UV-LDI MS data suggests that *ebony*’s effects on pigmentation and CHC length profiles are not linked at the level of the cuticle

Pigmentation synthesis in insect cuticles involves the secretion of biogenic amines (such as dopamine) by epidermal cells into the developing cuticle where they are oxidized into quinones that can form melanins or sclerotins that crosslink proteins (Figure 1A; reviewed in True, 2003 and Riedel *et al.*, 2011). To determine whether *ebony*’s effects on CHC length profiles depend on their function in pigmentation and sclerotization of the fly cuticle, we measured the relative abundance of individual CHCs in virgin females with different levels of pigmentation across the body. We crossed *pannier-GAL4* (Calleja *et al.* 2000) females with males from the *UAS-ebony-RNAi* effector line to generate flies with a dark, heavily melanized stripe down the dorsal midline (Figure 3A). We then used UV laser desorption/ionization mass spectrometry (UV-LDI MS) to take repeated measurements of CHCs along the thorax of females, targeting inside and outside the dark stripe (Figure 3A). Although we observed an upward trend in abundance from short to long CHCs, we did not detect a significant CHC lengthening effect like that observed between *ebony^CRISPR(1,2)^* flies and un-injected *vasa-Cas9* females (Figure 3B, Spearman’s ρ = 0.58, *P* = 0.13). Within the black cuticle, most CHCs detected by UV-LDI MS showed a decrease in abundance relative to brown cuticle (Figure 3B). This result suggests that *ebony* does not affect CHC length profiles through the pigmentation/sclerotization synthesis pathway, at least at the level of CHC/pigment deposition in the cuticle.

### 3.4 Abdominal pigmentation covaries with CHC length profiles in the *Drosophila* Genetic Reference Panel (DGRP)

The effects of *ebony* and *tan* mutants on CHC profiles described above suggest that variation in these genes might contribute to variation in both pigmentation and CHC profiles. Recently, Dembeck *et al.*(2015a,b) analyzed the genetic architecture of abdominal pigmentation and CHC composition in female *D. melanogaster* lines from the *Drosophila* Genetic Reference Panel (DGRP): Dembeck *et al.*(2015a) quantified abdominal pigmentation intensity in the 5^th^ and 6^th^ abdominal tergites (A5 and A6), and Dembeck *et al.* (2015b) investigated CHC profiles from the majority of the panel, but the relationship between the two traits was not examined. Using data from the 155 DGRP lines for which both pigmentation scores and CHC profiles were published, we tested the hypothesis that natural variation in pigmentation covaries with natural variation in CHC length profiles. In order to investigate CHC composition in a way that was comparable to the experiments described above, we divided the 155 DGRP lines into dark (N = 53), intermediate (N = 56), and light (N = 46) pigmentation groups using the 5^th^ abdominal tergite (A5) pigmentation scores (0–4) from Dembeck *et al* (2015a). Next, we tested whether females from dark, intermediate, or light pigmentation groups showed differences in their abundance of CHCs with different chain lengths relative to the 155 DGRP line average. We found that the group with the darkest A5 pigmentation showed lower levels of short chain CHCs and higher levels of long chain CHCs relative to the 155 line average (Figure 4A, Spearman’s ρ = 0.44, *P* < 0.01); the group with intermediate A5 pigmentation showed no relationship with CHC chain length (Figure 4B, Spearman’s ρ = 0.002, *P* = 0.98); and the group with lightest A5 pigmentation showed the opposite pattern as the dark group (Figure 4C, Spearman’s ρ = - 0.57, *P* = 1.0 x 10^-3^). We also compared CHC profiles in dark (N = 51), intermediate (N = 55), and light (N = 49) groups based on pigmentation of the 6^th^ abdominal tergite (A6), and found that, unexpectedly, the dark group did not show a significant CHC lengthening effect (Supplementary Figure S6A, Spearman’s ρ = 0.19, *P*= 0.25), and the intermediate group showed a CHC lengthening effect (Supplementary Figure S6B, Spearman’s ρ = 0.44, *P* < 0.01). However, the light group showed a significant CHC shortening effect as expected (Supplementary Figure S6C, Spearman’s ρ = −0.68, *P* < 1.0 x 10^-5^). These data suggest that darkly pigmented DGRP females show a pattern of CHC lengthening similar to the darkly pigmented loss-of-function *ebony^CRISPR(1,2)^* flies, and lightly pigmented DGRP females show a pattern of CHC shortening similar to lightly pigmented loss-of-function *tan^20A^* flies.

### 3.5 *ebony* and *tan* expression covaries with CHC length profiles in the DGRP

The DGRP genome-wide association (GWAS) study from Dembeck *et al.* (2015a) revealed that top variants associated with pigmentation are in *ebony, tan*, and *bab1*, consistent with variation in *ebony* expression level observed in the DGRP lines (Miyagi *et al.* 2015) and associations between pigmentation and these genes in studies of other *D. melanogaster* populations (Rebeiz *et al.* 2009a,b; Telonis-Scott *et al.* 2011; Takahashi and Takano-Shimizu 2011; Bastide *et al.* 2013; Endler *et al.* 2016; 2018). We therefore hypothesized that the differences in CHC length profiles seen in darkly and lightly pigmented DGRP females might be a consequence of expression variation at *ebony* and/or *tan*.

Using qRT-PCR, we quantified *ebony* and *tan* expression within 1 h after eclosion, which is when pigments determining adult body color are actively produced, in a sample of 23 DGRP lines that showed variable pigmentation. We then tested whether variation in *ebony* and *tan* expression covaried with CHC length profiles by categorizing the 23 DGRP lines into groups of low, intermediate, and high *ebony* or *tan* expression levels based on the qRT-PCR results, examining the average difference in individual CHC abundances between each expression group relative to the 23 line average, and plotting these values against CHC chain length (Figure 5).

**Figure 5.**
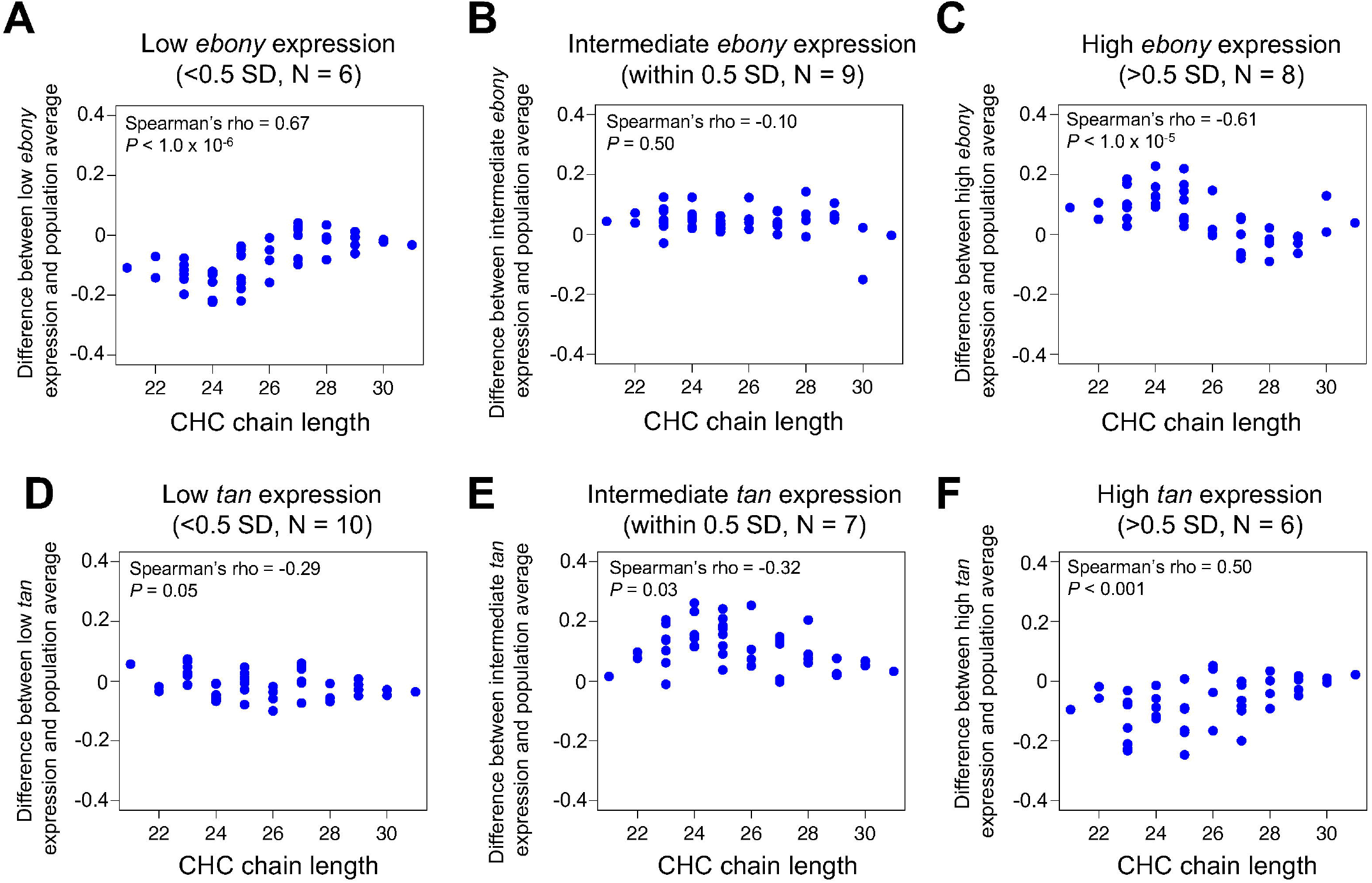
Variation in *ebony* and *tan* expression co-varies with CHC length profiles in the DGRP. CHC data was obtained from Dembeck *et al.* (2015b), and *ebony* and *tan* expression was quantified via qRT-PCR for 23 DGRP lines. **(A)** Difference in log-contrast of relative CHC intensity between DGRP females with low *ebony* expression and the 23 line average. **(B)** Difference in log-contrast of relative CHC intensity between DGRP females with intermediate *ebony* expression and the 23 line average. **(C)** Difference in log-contrast of relative CHC intensity between DGRP females with low *ebony*expression and the 23 line average. **(D)** Difference in log-contrast of relative CHC intensity between DGRP females with low *tan* expression and the 23 line average. **(E)** Difference in log-contrast of relative CHC intensity between DGRP females with intermediate *tan* expression and the 23 line average. **(F)** Difference in log-contrast of relative CHC intensity between DGRP females with high *tan* expression and the 23 line average.

Consistent with our hypothesis, the DGRP lines with low *ebony* expression showed lower levels of short chain CHCs, lines with high *ebony* expression showed higher levels of short chain CHCs, and lines with intermediate expression showed no change in CHC profiles (Figure 5A–C, Spearman’s ρ = 0.67, *P* < 1.0 x 10^-6^, Spearman’s ρ = −0.61, *P* < 1.0 x 10^-5^, Spearman’s ρ = −0.10, *P* = 0.50, respectively). Reciprocally, the DGRP lines with low or intermediate *tan* expression showed a slight increase in short chain CHCs, and lines with high *tan* expression showed a significant decrease in short chain CHCs (Figure 5D,F, Spearman’s ρ = −0.29, *P* = 0.05, Spearman’s ρ = −0.32, *P* = 0.03, Spearman’s ρ = 0.50, *P* < 0.001, respectively). Taken together, our results suggest that differences in *ebony* and *tan* gene expression have pleiotropic effects on both pigmentation and CHC length profiles that might cause these traits to covary in natural *D. melanogaster* populations.

## 4 Discussion

Pigmentation genes are often pleiotropic, with effects on vision, circadian rhythms, immunity, and mating behavior (reviewed in Wittkopp and Beldade, 2009; Takahashi, 2013). Here, we show that *ebony* and *tan* also affect CHC production, with the two genes altering CHC length profiles in opposing directions: *ebony^CRISPR(1,2)^* mutants had significantly higher levels of long chain CHCs, and *tan^20^* mutants had significantly higher levels of short chain CHCs. Our results suggest 1) that *ebony* and *tan* have a previously undescribed role in CHC synthesis and/or deposition and 2) that pleiotropy of both genes might influence the covariation of pigmentation and CHC composition.

### 4.1 Considering the pleiotropic effects of *ebony* and *tan* through changes in dopamine metabolism

Previous work has shown that changes in dopamine metabolism influence CHC composition in *Drosophila melanogaster*. Specifically, females homozygous for loss-of-function *Dopa-decarboxylase (Ddc)* temperature-sensitive alleles showed changes in CHC composition that could be reversed with dopamine feeding (Marican *et al.*, 2004; Wicker-Thomas and Hamann, 2008). Additionally, inhibiting dopamine synthesis by feeding wild-type females the tyrosine hydroxylase inhibitor L-AMPT altered CHC composition in a similar direction as the loss-of-function alleles (Marican *et al.*, 2004; Wicker-Thomas and Hamann, 2008). We found that feeding with L-AMPT affects CHC length composition, causing *ebony^CRISPR(1,2)^* and *ebony^CRISPR(3)^* mutants to have a more *tan^20^*-like CHC length profile (Figure 2A and Supplementary Figure S4). This result suggests that *ebony* and *tan* may affect CHC length composition through dopamine metabolism, but feeding *tan^20^* and wild-type females dopamine did not lead to CHC lengthening (Figure 2B,C and Supplementary Figure S5). Why did L-AMPT feeding affect CHC length composition while dopamine feeding did not? One possible reason is that L-AMPT is a potent inhibitor of tyrosine hydroxylase activity (Spector *et al.*, 1965), which processes tyrosine that flies ingest, whereas dopamine feeding might not cause significant changes in dopamine abundance in tissues relevant to CHC synthesis.

Another gene suggesting a possible link between CHC composition and dopamine is the *D. melanogaster apterous* gene. Loss of *apterous* gene function causes an increase in the proportion of long chain CHCs (Wicker and Jallon, 1995), and *apterous* mutants also show high levels of dopamine (Gruntenko *et al.*, 2003; Grutenko *et al.*, 2005; Grutenko *et al.*, 2012). These mutants also show low levels of juvenile hormone (JH) (Altaratz *et al.*, 1991), and treating decapitated females with methoprene to increase JH synthesis caused a decrease in long chain CHCs (Wicker and Jallon, 1995). The CHC lengthening and increased dopamine levels seen in *apterous* mutants resemble *ebony* mutants, but it is unknown whether *ebony* mutants show altered JH profiles. Further evidence supporting a role of JH and other ecdysteroids in determining CHC chain length comes from houseflies (Blomquist *et al.*, 1987). In *D. melanogaster*, ecdysteroid signaling was found to be required not only for CHC synthesis but also survival of the oenocyte cells that synthesize CHCs (Chiang *et al.*, 2016). An interesting future direction would be to test whether changes in dopamine metabolism in *ebony* or *tan* mutants influence CHC length composition through JH signaling. More broadly, a thorough genetic analysis focused on tissue-specific manipulation of dopamine is needed to deepen our understanding about its role in CHC synthesis.

### 4.2 CHC lengthening in *ebony* mutants does not seem to depend on changes at the level of the cuticle

Data from our tyrosine hydroxylase inhibition experiments supported the hypothesis that elevated dopamine levels in *ebony* mutants (as reported in Hodgetts and Konopka, 1973) affect CHC lengthening; however, it remains unclear which cells require *ebony* expression (and possibly dopamine metabolism) to influence CHC synthesis. We hypothesized that *ebony*-dependent changes of the fly cuticle itself might affect CHC deposition during fly development or CHC extraction in the laboratory, and found that all but one detected CHC showed an overall decrease in abundance in dark cuticle relative to light cuticle. However, we note that these differences might be due to changes in the physical properties of dark versus light cuticle as they interact with the UV-LDI instrument. We also note that *ebony^CRISPR(3)^* and *ebony^CISPR(4)^* mutants had darkly pigmented cuticle like *ebony^CRISPR(1,2)^* mutants but CHC length profiles similar to wild-type flies, suggesting that *ebony* and tan’s effects on CHC length composition can be separated from their role in pigmentation synthesis. For example, *ebony* expression in glia is necessary for normal circadian rhythms in *D. melanogaster* but not pigmentation (Suh and Jackson, 2007). We tested whether knocking down *ebony* in oenocytes affected CHC length composition and found that it did not, thus the specific cells required for *ebony* and *tan*’s effects on CHC synthesis remain unknown.

### 4.3 Patterns of CHC composition and pigmentation along clines in natural populations

Identifying the pleiotropic effects of *ebony* and *tan* on pigmentation and CHCs is important because it suggests that these genes might contribute to the covariation of both traits in natural populations. For example, selection for *ebony*- or *tan*-dependent pigmentation variation might also cause variation in CHC length composition without selection acting directly on this trait. Alternatively, selection for long chain CHCs with higher melting temperatures (Gibbs and Pomonis, 1995; Gibbs, 1998) in drier climates might cause a correlated increase in pigmentation intensity. Indeed, we found that variation in abdominal pigmentation covaries with both *ebony* and *tan* gene expression as well as CHC length profiles in directions predicted by *ebony* and *tan* mutants among the DGRP lines, which were derived from flies isolated from a single, natural population (Ayroles *et al.* 2009; Mackay *et al.* 2012; Huang *et al.* 2014). However, this finding does not necessarily imply variation in both traits is caused by the same gene(s) nor that these traits will always co-evolve; for example, individuals with dark pigmentation may coincidentally possess alleles that are in linkage disequilibrium that cause a CHC lengthening phenotype. Comparing the phenotypic frequency of pigmentation and CHC length composition phenotypes within and between the same populations that are undergoing adaptation to common environments will help answer this question. In Africa, for example, *D. melanogaster* populations repeatedly show a strong positive correlation between elevation and dark pigmentation, suggesting that environments at high altitudes might select for darkly pigmented flies (or some other trait that correlates with pigmentation) (Pool and Aquadro, 2007; Bastide *et al.*, 2014). It will be interesting to know whether these populations also show an increase in abundance of long chain CHCs.

Both pigmentation and CHC length profiles vary along altitudinal and latitudinal clines in natural *Drosophila* populations, suggesting that ecological factors such as humidity or temperature play a role in shaping variation in at least one of these traits. At higher altitudes or latitudes, populations often showed darker pigmentation profiles in Europe, India, and Australia (Heed and Krishnamurthy, 1959; David *et al.*, 1985; Capy *et al.*, 1988; Das, 1995; Munjal *et al.*, 1997; Parkash and Munjal, 1999; Pool and Aquadro, 2007; Telonis-Scott *et al.*, 2011; Parkash *et al.*, 2008a; Parkash *et al.*, 2008b; Matute and Harris, 2013). In Africa, however, latitude and pigmentation intensity showed a negative correlation, so this relationship is not universal (Bastide *et al.*, 2014). For CHCs, Rajpurohit *et al.* (2017) reported that *D. melanogaster* populations at higher latitudes showed more short chain CHCs, whereas populations at lower latitudes showed more long chain CHCs in the United States. Frentiu and Chenoweth (2010) similarly found that populations at high latitudes along a cline in Australia showed more short chain CHCs and fewer long chain CHCs. These patterns do not match predictions based on the pleiotropy we observed: flies at higher latitudes tend to have darker pigmentation and higher levels of short chain CHCs whereas *ebony^CRISPR(1,2)^* mutants, for example, have darker pigmentation and lower levels of short chain CHCs. To the best of our knowledge, pigmentation (nor *ebony* or *tan* expression) and CHC length composition have not been simultaneously measured in flies from the same cline, making it difficult to discern whether pigmentation and CHC composition covary in the wild in ways predicted by the mutant data. For example, Frentiu and Chenoweth (2010) measured CHCs from populations along the east coast of Australia, but they did not include populations from higher latitude coastal regions with darker pigmentation and lower *ebony* expression in newly eclosed adults (Telonis-Scott *et al.* 2011). Comparing variation in both traits within and between populations along latitudinal and/or altitudinal clines will make it clearer if and to what extent pigmentation and CHC composition covary and whether variation in these features is accompanied by changes in *ebony* and *tan* expression.

## 6 Author Contributions

J.H.M., N.A., P.J.W., J.Y.Y., and A.T. conceived the project; J.H.M., N.A., T.B., K.D., and J.Y.Y. collected the data; J.H.M, N.A., T.B., K.D., J.Y.Y., and A.T. analyzed the data; and J.H.M., P.J.W., J.Y.Y., and A.T wrote the paper.

## 8 Acknowledgments

We thank members of the Takahashi, Wittkopp, and Yew labs and Aki Ejima for helpful discussions; John True, Stephen Goodwin, Scott Pletcher, Rainbow Transgenics Inc., the Bloomington Drosophila Stock Center, and the Vienna Drosophila RNAi Center for fly stocks; and Rainbow Transgenics Inc., for fly injections.

## 10 Data Availability Statement

The raw data supporting the conclusions of this manuscript will be made available by the authors, without undue reservation, to any qualified researcher.

