## Supplementary material for "Pleiotropic effects of *ebony* and *tan* on pigmentation and cuticular hydrocarbon composition in *Drosophila melanogaster*": Table S1

**Table 1. Common CHCs in ♀ *D. melanogaster***

| Category |  | Elemental  Formula | | Common Notation^†^ |
| --- | --- | --- | --- | --- |
| alkane |  | C21 H44 |  | nC21 |
| monoene |  | C22 H44 |  | C22:1 |
| alkane |  | C22 H44 |  | nC22 |
| methyl branched | | C22 H46 |  | 23Br |
| diene |  | C23 H44 |  | 7.11TD |
| monoene |  | C23 H46 |  | 9-T |
| monoene |  | C23 H46 |  | 7-T |
| monoene |  | C23 H46 |  | 5-T |
| alkane |  | C23 H46 |  | nC23 |
| alkane |  | C24 H50 |  | nC24 |
| methyl branched | | C24 H50 |  | 25Br |
| diene |  | C25 H48 |  | 7.11PD |
| monoene |  | C25 H50 |  | 9-P |
| monoene |  | C25 H50 |  | 7-P |
| alkane |  | C25 H52 |  | nC25 |
| alkane |  | C26 H54 |  | internal standard |
| methyl branched | | C26 H54 |  | 27Br |
| diene |  | C27 H52 |  | 7.11HD |
| monoene |  | C27 H54 |  | 9-H |
| monoene |  | C27 H54 |  | 7-H |
| alkane |  | C27 H56 |  | nC27 |
| methyl branched | | C28 H58 |  | 29Br |
| diene |  | C29 H56 |  | 7.11ND |
| alkane |  | C29 H60 |  | nC29 |

^†^Br: methyl branched; T: tricosene; P: pentacosene; H: heptacosene; TD: tricosadiene;

PD: pentacosadiene; ND: nonacosadiene
