## Supplementary figures and images for "Pleiotropic effects of *ebony* and *tan* on pigmentation and cuticular hydrocarbon composition in *Drosophila melanogaster*"

### Figure S1

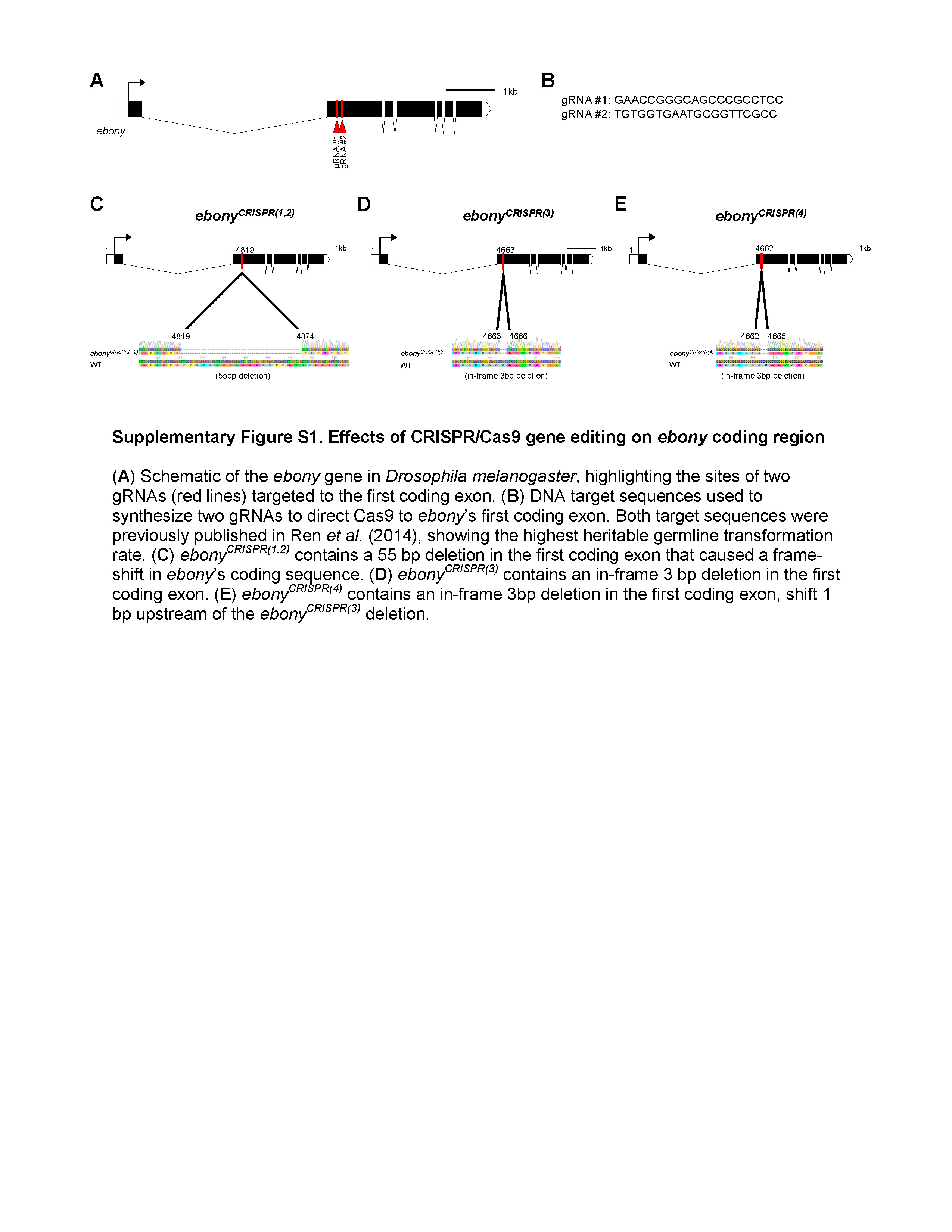

### Figure S2

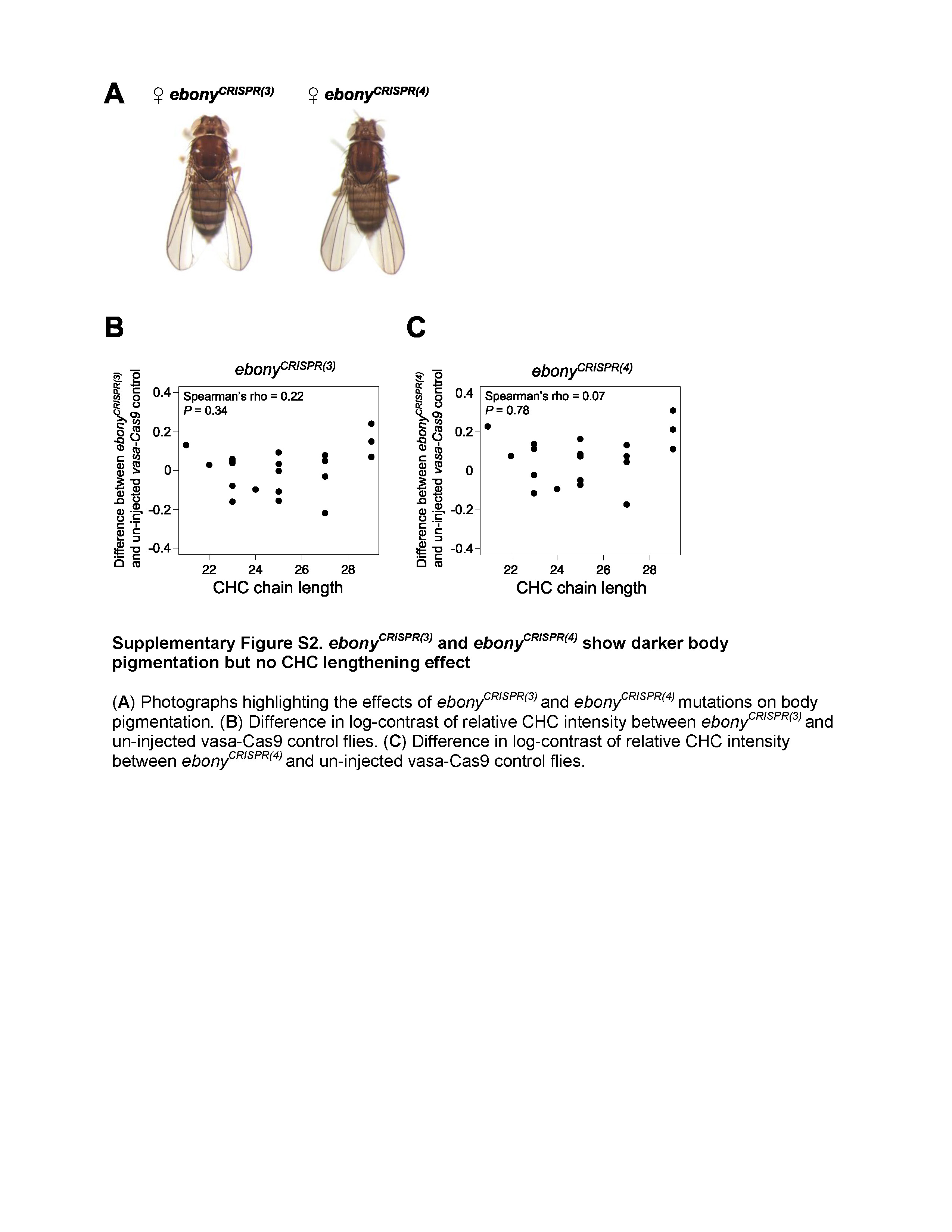

### Figure S3

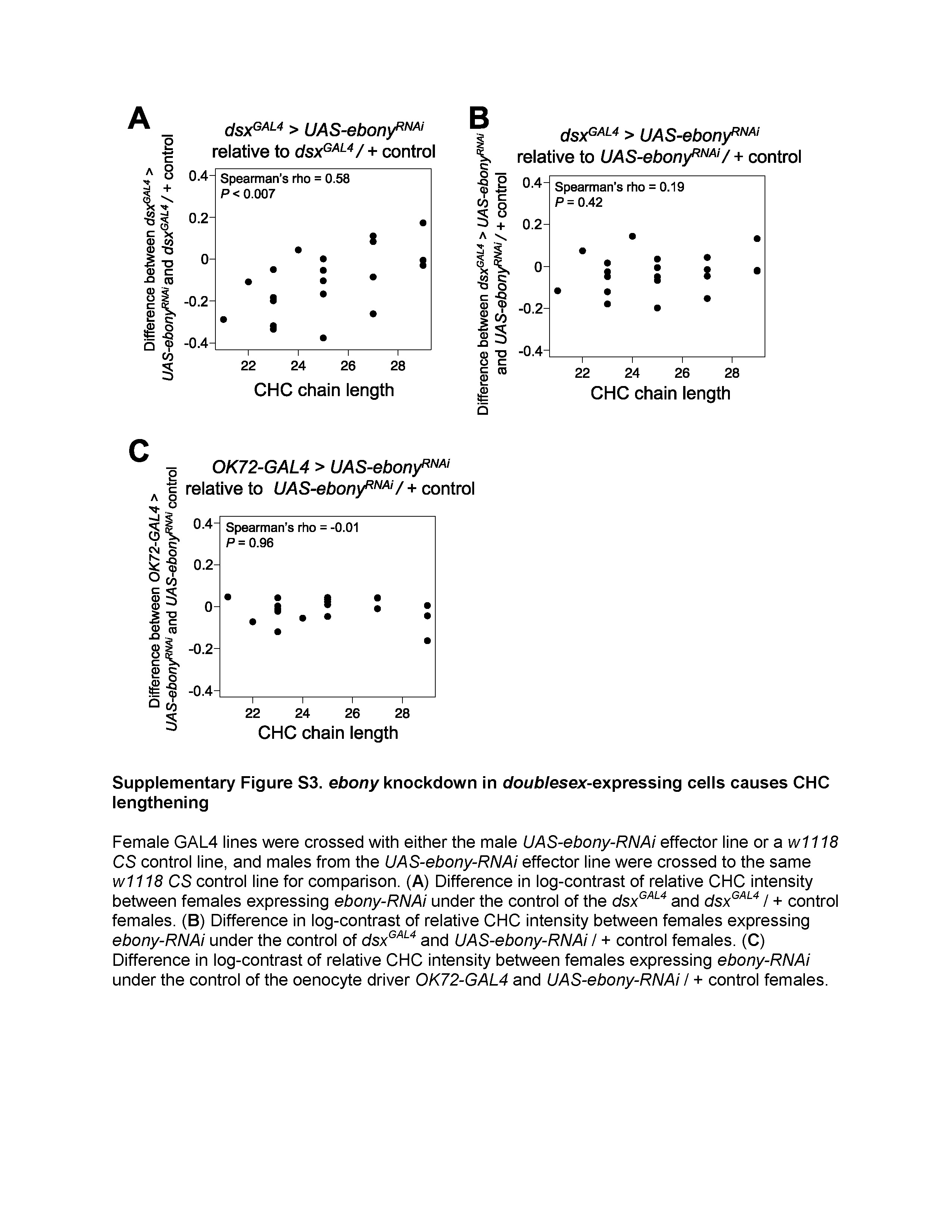

### Figure S4

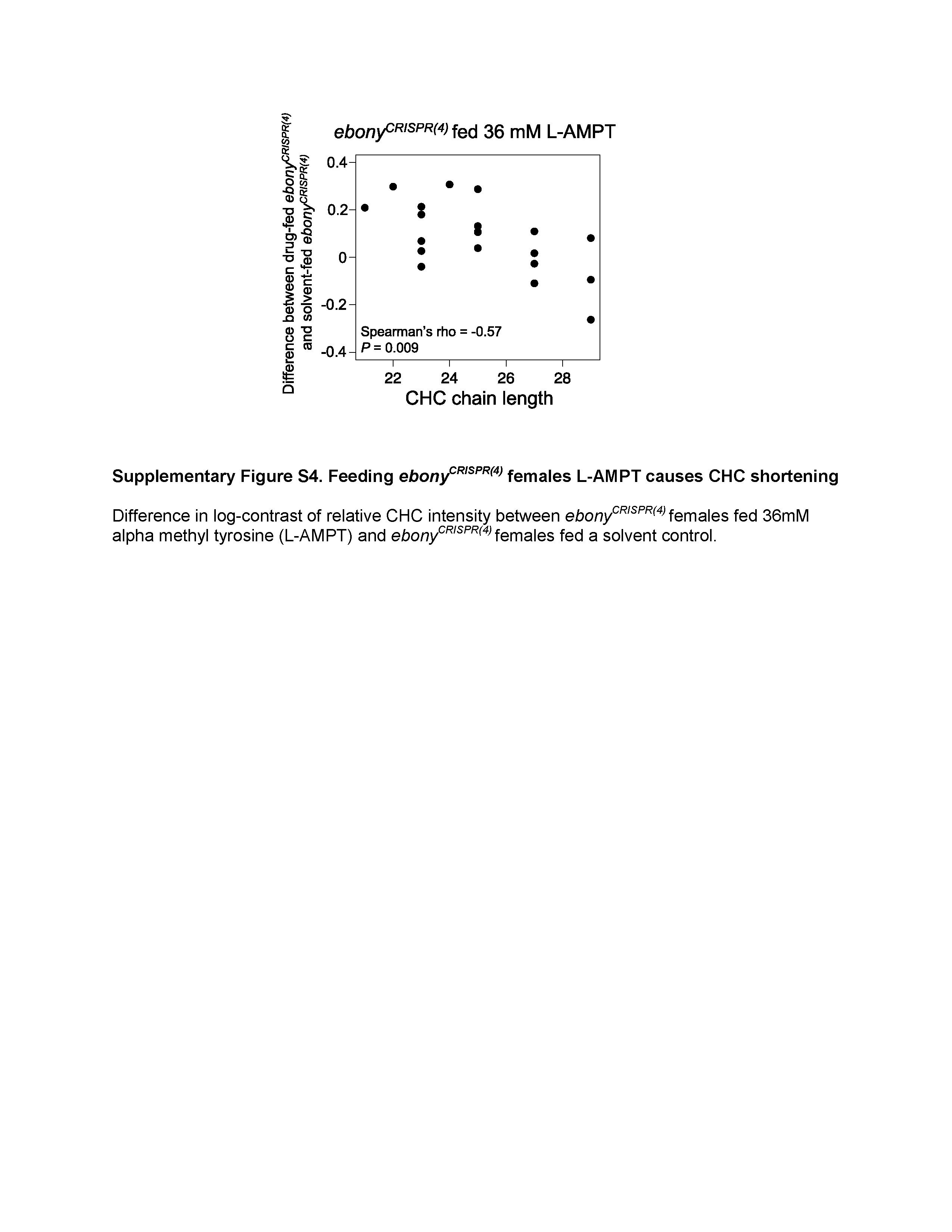

### Figure S5

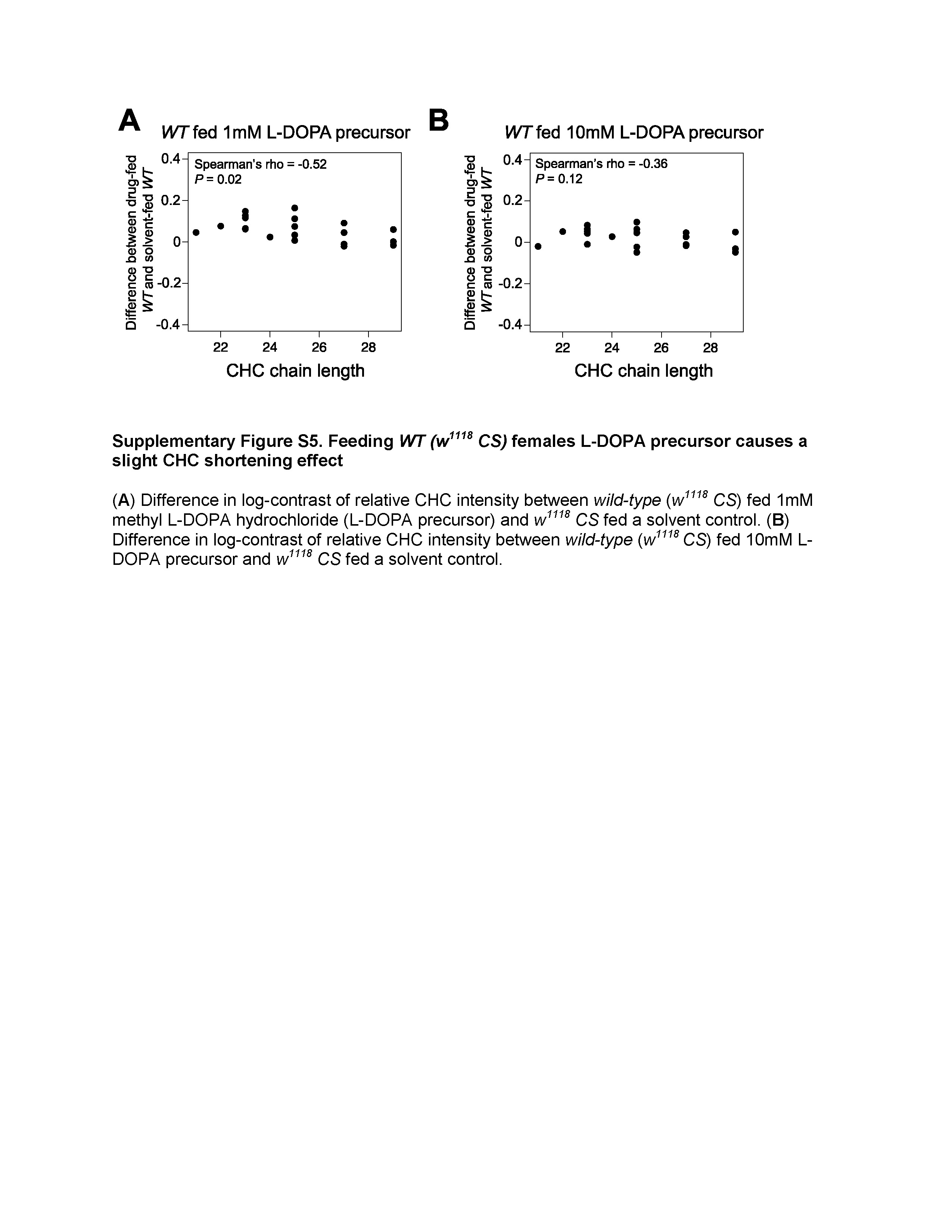

### Figure S6

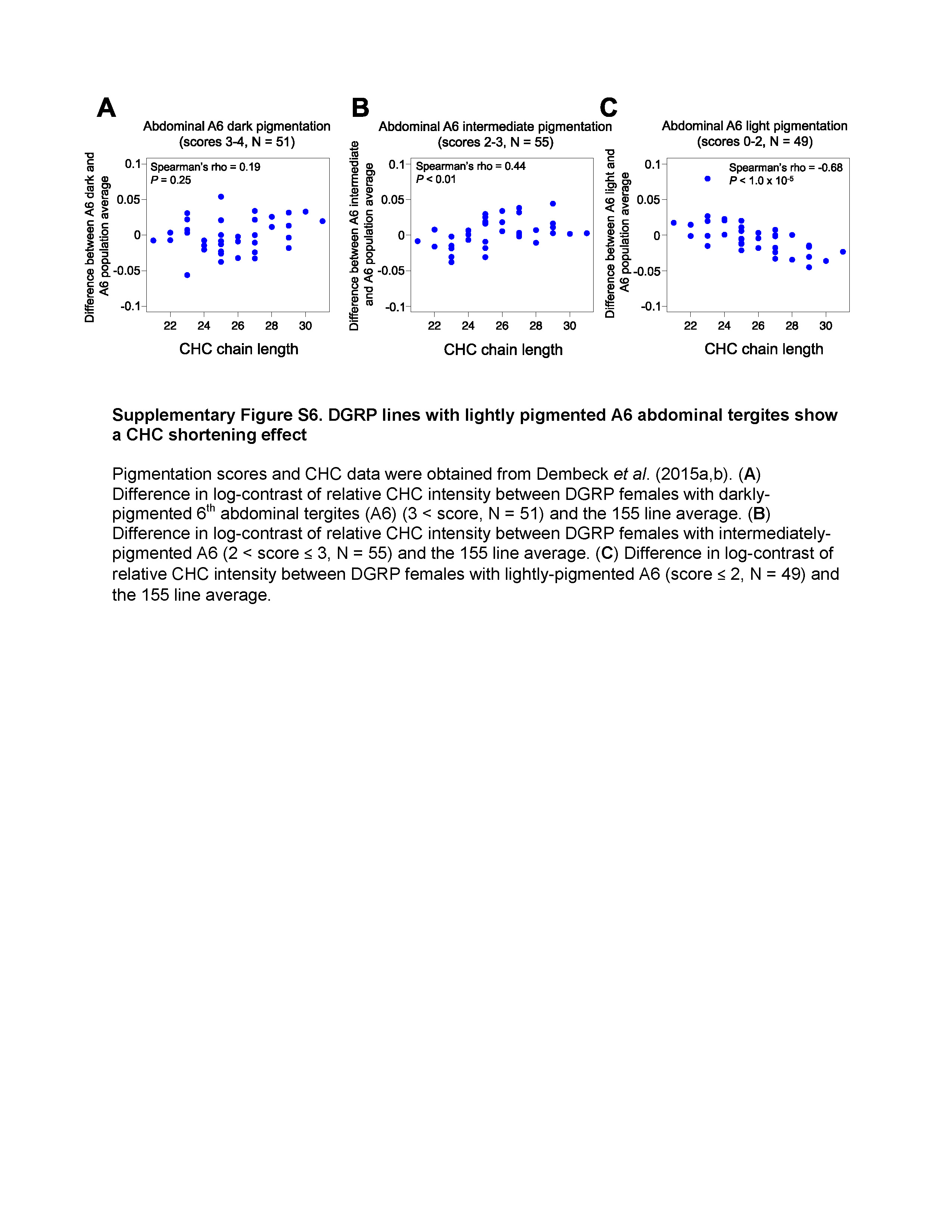
